# Modelling human KCNT1-epilepsy in *Drosophila*: a seizure phenotype and drug responses

**DOI:** 10.1101/2023.04.11.536495

**Authors:** Rashid Hussain, Chiao Xin Lim, Zeeshan Shaukat, Anowarul Islam, Emily A. Caseley, Jonathan D. Lippiat, Grigori Y. Rychkov, Michael G. Ricos, Leanne M. Dibbens

## Abstract

Mutations in the *KCNT1* potassium channel cause severe forms of epilepsy which are resistant to current treatments. *In vitro* studies have shown that *KCNT1-*epilepsy mutations are gain of function, significantly increasing K^+^ current amplitudes. To investigate if *Drosophila* can be used to model human *KCNT1* epilepsy, we generated *Drosophila melanogaster* lines carrying human *KCNT1* with the patient mutation G288S, R398Q or R928C. Expression of each mutant channel in GABAergic neurons gave a seizure phenotype which was sensitive to drugs currently used to treat patients with *KCNT1*-epilepsy. Cannabidiol showed the greatest reduction of the seizure phenotype while some drugs increased the seizure phenotype. Our study shows that *Drosophila* can be used to model human *KCNT1*-epilepsy and potentially used as a tool to assess new treatments for *KCNT1* epilepsy.

## Introduction

Mutations in *KCNT1* have been identified in a range of epilepsies with drug-resistant seizures (Lim, Ricos, Dibbens, & Heron, 2016). *KCNT1* mutations are the major cause of epilepsy of infancy with migrating focal seizures (EIMFS) (Barcia et al., 2012; Bonardi et al., 2021; Lim et al., 2016), where cognitive and developmental regression follow seizure onset. *KCNT1* mutations also cause other severe epilepsies beginning in infancy, including West Syndrome and Otahara Syndrome (Lim et al., 2016), as well as a range of focal epilepsies, which can have a later age of onset (in childhood or adolescence), including sleep-related hypermotor epilepsy (Heron et al., 2012; Lim et al., 2016; Moller et al., 2015). *KCNT1*-epilepsy is often debilitating and can include developmental regression and cognitive impairment after seizure onset (Bonardi et al 2021).

*KCNT1* encodes an ion channel which is a major contributor of the sodium-activated potassium IK_Na_ current which down regulates neuronal excitability. Following a rise in intracellular [Na^+^], KCNT1 channels are thought to increase K^+^ current and prolong the slow afterhyperpolarisation phase following an action potential, thereby reducing the chance of repetitive neuronal firing (Fisher et al., 2005; Köhling & Wolfart, 2016; Lim et al., 2016).

Almost all *KCNT1* epilepsy mutations are heterozygous and missense, predicted to change a single amino acid in the protein (Bonardi et al., 2021) and *de novo* mutations most often account for severe cases. All mutations analysed *in vitro* to date (apart from T314A, Rychkov et al 2022) significantly increase K^+^ currents in comparison to normal KCNT1 channels (Kim et al., 2014; Milligan et al., 2014; Rizzo et al., 2016; Rychkov et al., 2022; Tang et al., 2016). There is evidence suggesting that increased K^+^ currents in inhibitory neurons may be associated with *KCNT1*-related seizures (Gertler, Cherian, DeKeyser, Kearney, & George, 2022; Rychkov et al., 2022; Shore et al., 2020). Reducing the activity of inhibitory neurons may explain how neuronal hyperexcitability, the mechanism underlying seizures, occurs in *KCNT1*-epilepsy.

There are currently no effective treatments for *KCNT1*-epilepsy and no drugs available that specifically target the KCNT1 channel. Demonstration that increased K^+^ current due to overactivity of the KCNT1 channel is associated with seizures suggests that reducing or blocking KCNT1 channel activity may inhibit seizures, and this has directed new drug screening efforts (Cole et al., 2020). Currently, anti-epileptic drugs including carbamazepine, vigabatrin and valproic acid, as well as the drug cannabidiol (CBD) are used as treatments for *KCNT1-*epilepsy (Fitzgerald et al., 2019), (Bonardi et al., 2021; Borlot et al., 2020; Poisson, Wong, Lee, & Cilio, 2020). However, even combinations of multiple drugs (sometimes eight or more) usually fail to suppress seizures, with some patients continuing to experience up to one hundred seizures per day (Bonardi et al., 2021). *In vitro* studies have shown that the ion channel blocker quinidine reduces the increased current amplitude produced by *KCNT1*-epilepsy mutations (Milligan et al., 2014; Mikati et al., 2015; Rizzo et al., 2016). However, it has shown variable results in patients and can have serious side effects (Bearden et al,. 2014, Mikati et al., 2015, Chong et al,. 2016, Bonardi et al., 2021).

In this study we sought to investigate if *Drosophila* could be used to model *KCNT1*-epilepsy and if the seizure phenotype responded to some of the drugs currently used to treat patients. Many of the genes and pathways identified in human epilepsy are highly conserved in *Drosophila* (Baines, Giachello, & Lin, 2017). The animal has been used to model other forms of human epilepsy, including Dravet Syndrome and Generalised Epilepsy with Febrile Seizures Plus (GEFS+) due to mutations in the sodium channel gene *SCN1A* (Parker, Padilla, Du, Dong, & Tanouye, 2011; Takai et al., 2020) and can make powerful tools for therapeutic screens (Dare et al., 2020). The majority of *KCNT1*-epilepsy mutations cluster around three, highly conserved, functional domains in the KCNT1 protein (Lim et al., 2016). A mutation from each region was selected to be analysed: c.862G>A, p.G288S in the S5 segment of the pore domain, c.1193G>A, p. R398Q in the RCK1 (Regulator of K^+^ Conductance) domain and c.2782C>T, p.R928C adjacent to the NAD^+^ binding domain (Figure 1). G288S and R398Q mutations have been idenitfied in patients with a range of epilepsy phenotypes, ranging from very severe infantile onset epilepsies to less severe and later onset focal epilepsies, whereas R928C mutations are only associated with the latter. The three mutations are recurrently found in patients, collectively accounting for approximtaely 20% of cases (Bonardi et al., 2021; Lim et al., 2016). Thus, successful modelling of human *KCNT1*-epilepsy with these mutations would be relevant for a significant proportion of patients.

**Fig. 1.**
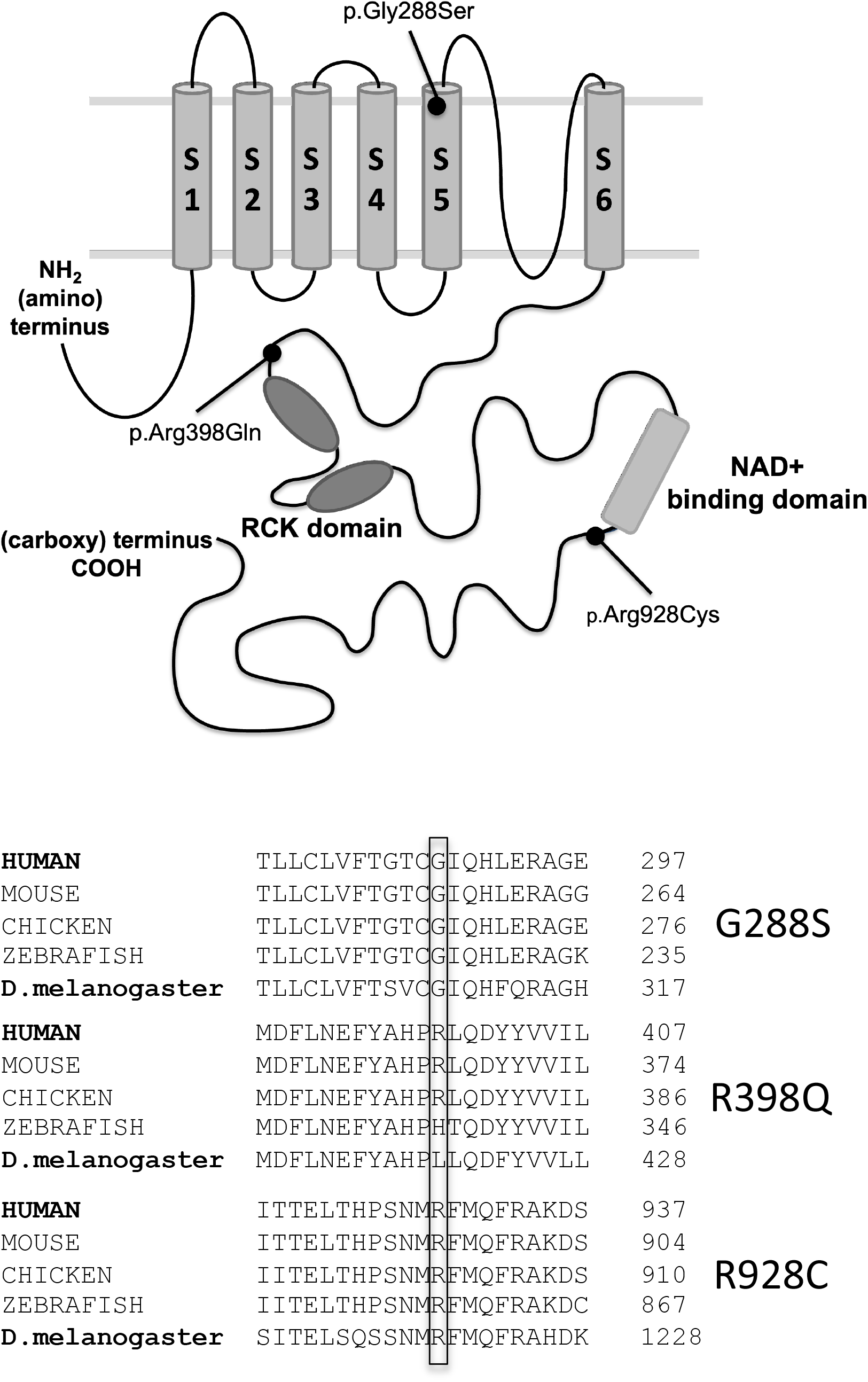
Position and evolutionary conservation of mutated amino acid residues. Position and evolutionary conservation of the amino acid residues altered by the three patient KCNT1 missense mutations investigated in this study, G288S, R398Q and R928C. **a**, Schematic diagram of the KCNT1 channel showing the positions of the three mutations investigated in this study. **b**, Alignment of the orthologous KCNT1 proteins found in different species showing the high evolutionary conservation of the amino acids altered by the 3 KCNT1 mutations investigated in this study.

To generate the *Drosophila* models, human *KCNT1* with the patient mutation G288S, R398Q or R928C was introduced into *Drosophila* by transgenesis. As in humans, the major inhibitory neurotransmitter in the *Drosophila* central nervous system is γ-aminobutyric acid (GABA) (Wilson & Laurent, 2005). Previous studies have suggested that reduced activity of inhibitory GABAergic neurons may be associated with seizures (Shore et al 2020, Gertler et al 2022, Rychkov et al 2022). The mutant KCNT1 channels were expressed in different neural tissues, including GABAergic neurons, and investigated for a seizure phenotype using bang sensitive behavioural assays (Lee & Wu, 2002; Song & Tanouye, 2008). To investigate the potential of the *Drosophila* models in assessing treatments for patients with *KCNT1* epilepsy, some of the drugs currently used to treat people with KCNT1 epilepsy were analysed for their effects on the seizure phenotype in *Drosophila*.

## Results

### Investigation of a seizure phenotype in Drosophila with G288S, R398Q or R928C mutant KCNT1

To investigate if *Drosophila c*arrying *KCNT1* mutations showed a seizure phenotype, three transgenic lines were generated with mutated human *KCNT1*. (Figure 1a,b). Each *Drosophila* line carried a human *KCNT1* transgene with a heterozygous missense mutation that has been identified in patients, G288S, R398Q or R928C. Wild type (normal) human *KCNT1* (NM_020822.3) was used as a control. The UAS-GAL4 expression system (Brand & Perrimon, 1993) was used to drive expression of the human *KCNT1* transgene. Each *Drosophila* line contained an upstream activating sequence (UAS) positioned before the *KCNT1* transgene. Genetic crosses were used to introduce the GAL4 transcription factor under the control of selected promoters, to drive expression of the wild type or mutant KCNT1 channels in different tissues. The pan-neural promoter *elav*^*C155*^*-GAL4* was used to drive *KCNT1* expression in all neurons. Expression of wild type human *KCNT1* showed no effect on viability, while expression of G288S, R398Q and R928C mutant *KCNT1* caused juvenile lethality at the embryonic stage, thus a seizure phenotype was not able to be investigated (Table 1).

**Table 1.** Effect of wild type (WT) and mutant human KCNT1 transgene expression in neuronal subsets.

| Driver<br>Mutant | **Pan-neuronal**<br>**Elav-GAL4** | **Cholinergic**<br>**Chat-Gal4** | **GABAergic**<br>**GAD1-GAL4** | **Glial**<br>**Repo-GAL4** |
| --- | --- | --- | --- | --- |
| **UAS-KCNT1-WT** | Viable no effect | Viable no effect | Viable no effect | Viable no effect |
| **UAS-KCNT1-G288S** | Embryonic<br>Lethal | Embryonic<br>Lethal | Viable- Bang<br>Sensitive | Embryonic<br>Lethal |
| **UAS-KCNT1-R398Q** | Embryonic<br>lethal | Embryonic<br>Lethal | Viable- Bang<br>Sensitive | Embryonic<br>Lethal |
| **UAS-KCNT1-R928C** | Embryonic<br>lethal | Embryonic<br>Lethal | Viable- Bang<br>Sensitive | Embryonic<br>Lethal |

Expression of wild type and mutant *KCNT1* was driven in GABAergic neurons using the GAD1 promoter (Ng et al., 2002). This gave surviving adults which were investigated for a seizure phenotype in bang-sensitive behavioural assays (Lee & Wu, 2002; Song & Tanouye, 2008) and the percentage of animals showing a seizure phenotype were calculated for each genotype. The mutant *KCNT1* lines G288S, R398Q and R928C each showed a seizure phenotype, while expression of wild type human *KCNT1* in GABAergic neurons did not (Figure 2). R398Q gave the strongest seizure phenotype with 48% of animals showing seizure activity followed by G288S with 41% and R928C with 38%.

**Figure 2.**
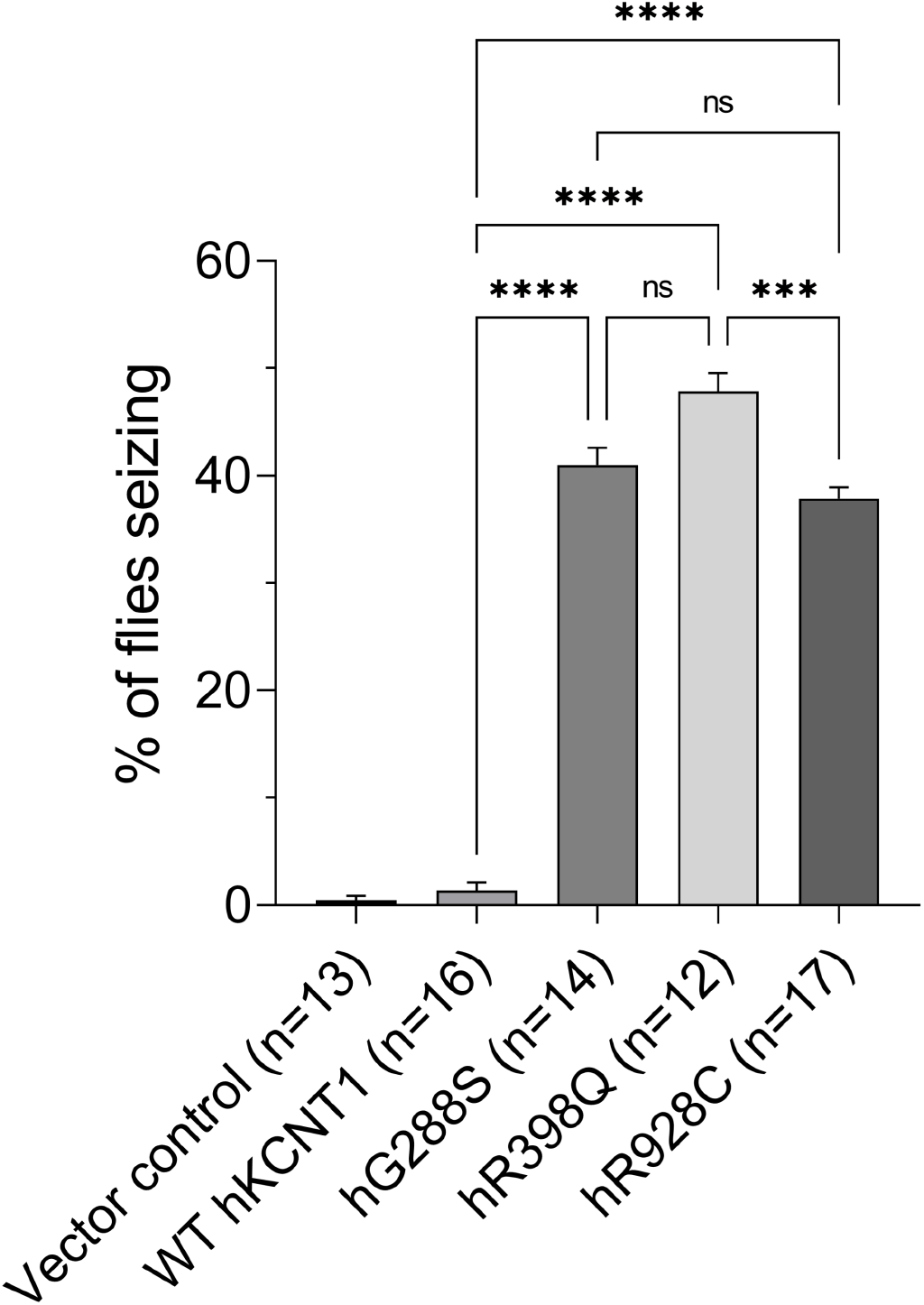
Expressing Human KCNT1 mutants in GABAergic neurons of *Drosophila* give rise to seizures in a bang sensitive behavioural assay. *Drosophila* with the vector control and *Drosophila* expressing either wild type (WT) or G288S, R398Q or R928C mutant human *KCNT1* in GABAergic neurons were analysed in the bang sensitive behavoural seizure assay. Percentage of *Drosophila* showing a seizure phenotype are shown for each line. **** p<0.0001, *** p<0.001, ns - no significant difference

The *KCNT1* transgenes were then put under the control of the *CHAT-GAL4* driver containing the *Choline-Acetyltransferase* (*Chat*) gene promoter to drive expression in excitatory cholinergic neurons (Salvaterra & Kitamoto, 2001). Expression of wild type *KCNT1* in cholinergic neurons gave adult flies which did not show a seizure phenotype (data not shown), while expression of the three mutant *KCNT1* lines was embryonic lethal (Table 1). Expression of wild type *KCNT1* mutant in glia, using the *reversed polarity* (*Repo)-GAL4* driver, did not give a seizure phenotype in bang-sentitive assays, and expression of the three mutant *KCNT1* lines G288S, R398Q and R928C was embryonic lethal (Table 1).

### In vitro effects of currently used drugs on KCNT1 channels

Having found that expression of G228S, R398Q and R928C mutant *KCNT1* in GABAergic neurons gives a seizure phenotype, we next investigated if the phenotype generated by each of the three mutations responded to some of the drugs most commonly used to treat patients with *KCNT1*-epilepsy. In the initial experiments, we looked at the *in vitro* effects of the drugs on KCNT1 channels in human cells using a HEK293T cell expression system and patch clamping analysis. This provided a baseline for the effects of the drugs directly on the KCNT1 channels. We analysed the effects of of carbamazepine, valproic acid, vigabatrin and cannabidiol (CBD). The *in vitro* effects of quinidine were not investigated as part of this study as these have been published previously (Milligan et al., 2014; Rizzo et al., 2016). Each drug was first tested on wild type KCNT1 channels at a range of concentrations, including well above their currently accepted potential doses for therapeutic use. The drugs were applied to the bath through the perfusion system and the amplitude of wild type KCNT1 current was monitored by applying voltage ramps between - 120 and 120 mV every 2 seconds. Carbamazepine (100 μM), valproic acid (30 μM), and vigabatrin (50 μM) showed no effect on the current amplitude produced by wild type KCNT1 within 5-10 min of application (Figure 4a). Since the drugs showed no effects on the wild type KCNT1 channels we did not test their *in vitro* effects on the mutant channels. In contrast, CBD (25 μM) inhibited almost 90% of the wild type KCNT1 current in HEK293T cells (Figure 4a).

Consistent with previous findings, the amplitudes of KCNT1 currents in cells expressing the G288S, R398Q and R928C mutants were significantly larger than those expressing wild type KCNT1 (Figure 3). Addition of CBD to HEK293T cells expressing each of the KCNT1 mutant channels showed a significant reduction in K^+^ current amplitude (Figure 4B). CBD was seen to inhibit KCNT1 mutant currents with higher potency compared to wild type KCNT1 (Figure 4b). The IC_50_ for wild type KCNT1 was approximately 10 times higher than for the mutant channels (Figure 4b). The time course of inhibition was similar between wild type and mutant KCNT1 channels (Figure 4c, data for wild type and R928C KCNT1 is shown), and CBD was fully washable.

**Figure 3.**
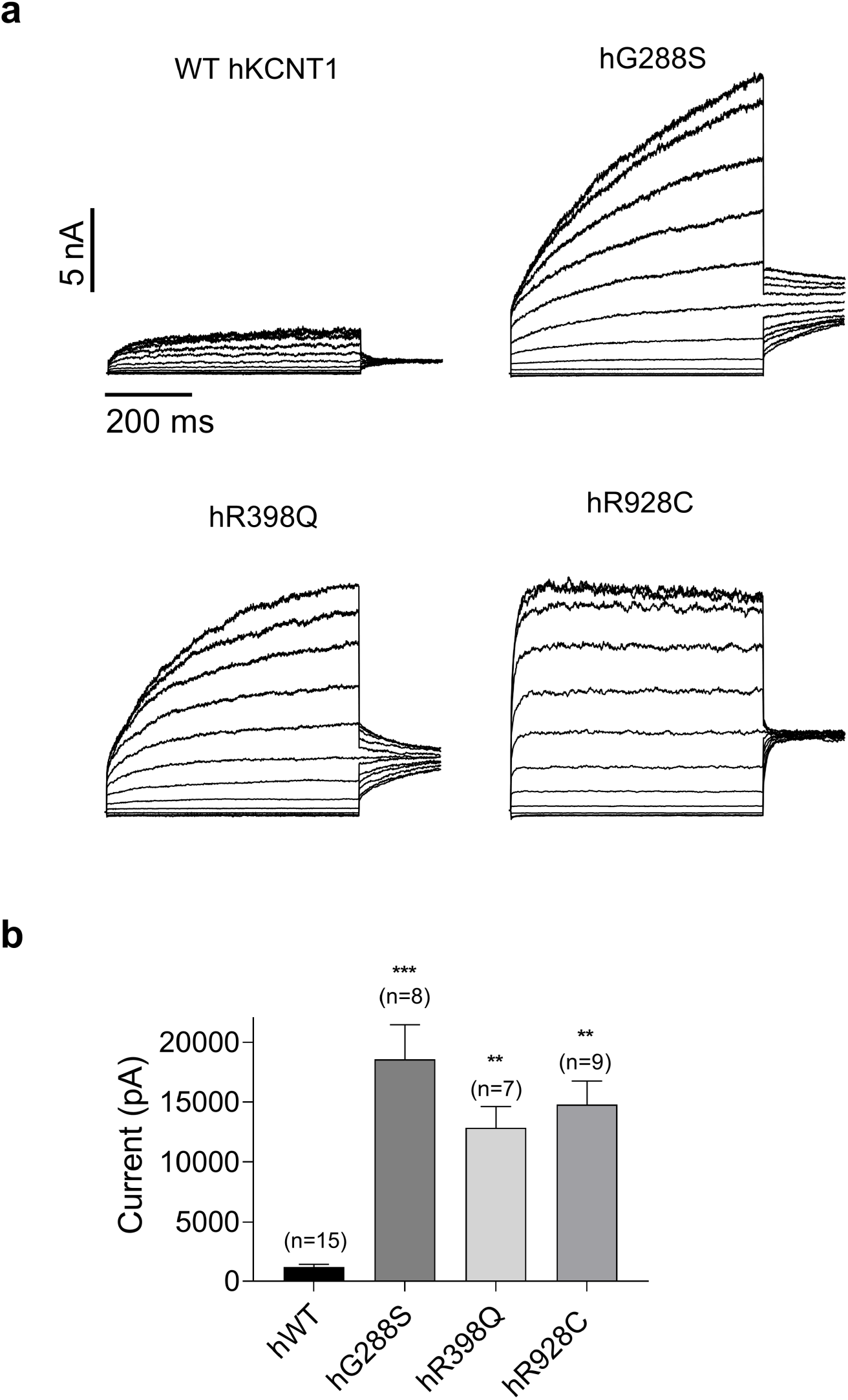
WT and Mutant KCNT1 currents recorded in HEK293T cells. **a**. KCNT1 currents recorded in HEK293T cells in response to the voltage steps ranging from -120 mV to 80 mV in 20 mV increments followed by a voltage step to 0 mV. **b**. The average amplitudes of the WT and mutant KCNT1 currents at 10 mV obtained from the currents recorded in response to the voltage ramps between -120 and 120 mV

Investigation of KCNT1 single channel activity using inside-out patches showed that CBD applied to the intracellular side of the membrane inhibited both wild type and mutant KCNT1 channels at lower concentrations, compared to the extracellular applications (Figure 5), suggesting intracellular CBD binding site on the KCNT1 protein. Furthermore, single channel data indicated that CBD blocked whole-cell KCNT1 currents by decreasing the open probability of the channel without affecting single channel conductance (Figure 5).

**Figure 4.**
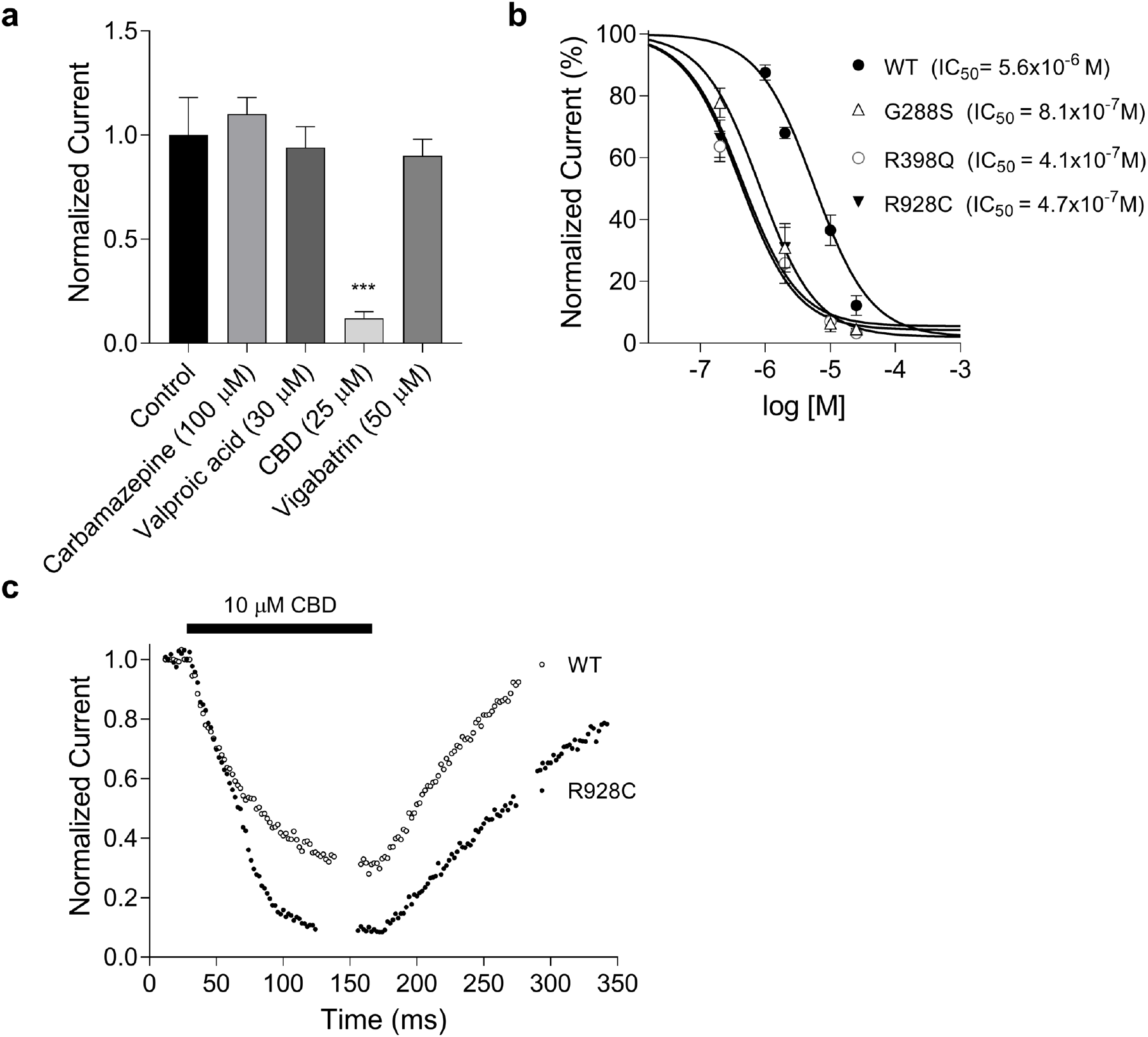
The *in vitro* effects of drugs on KCNT1 channels. **a**. The average normalised WT KCNT1 current amplitude recorded at 10 mV in the presence of the drugs. **b**. The dose-response curves of CBD inhibition of the wild type (WT) and mutant KCNT1 channels. **c**. The time course of KCNT1 inhibition by 10 μM CBD followed by the washout.

**Figure 5.**
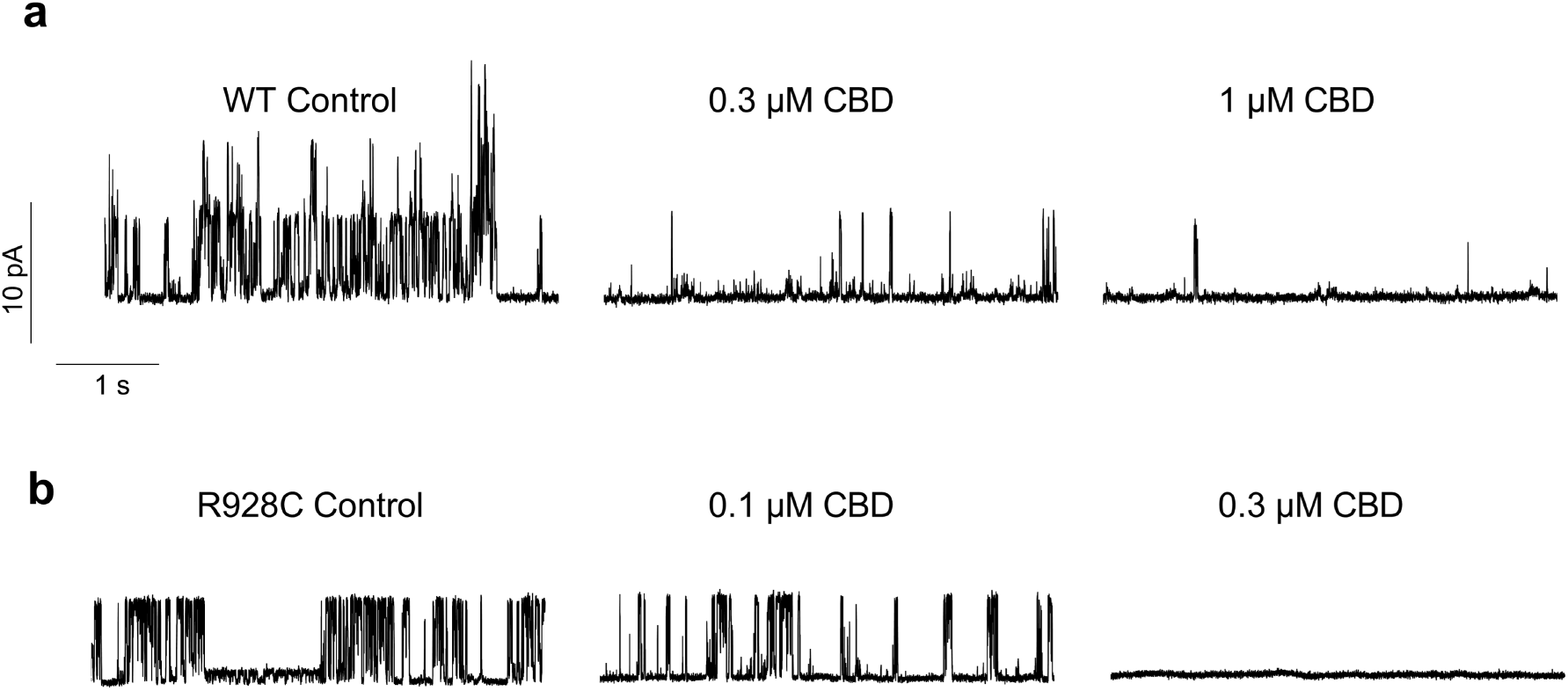
The effect of CBD on single WT and mutant (R928C) KCNT1 channels. Single WT (**a**) and mutant channels (**b**) were recorded in inside-out patches under control conditions and in the presence of the indicated ammounts of CBD in the bath.

### Responses of KCNT1 G228S, R938Q and R928C Drosophila seizure models to currently used drug treatments

We next investigated the effects of the drugs carbamazepine, valproic acid, vigabatrin, quinidine and CBD on the seizure phenotype of the three *Drosophila* mutant *KCNT1* lines. We used feeding experiments with a range of drug concentrations, followed by the bang sensitive behavioural assay. None of the drugs were seen to have an effect on the viability or behaviour of *Drosophila* expressing wild type KCNT1 (data not shown).

Carbamazepine, valproic acid and quinidine, were each seen to exacerbate the seizure phenotype in the three *KCNT1* mutant *Drosophila* lines (Figure 6). Vigabatrin was seen to reduce the seizure phenotype in the lines expressing G288S and R398C mutant KCNT1, while it increased the seizure phenotype in the R928C mutant. CBD showed significant reduction of the seizure phenotype in all three *KCNT1* mutant lines and showed a dose-dependent response (Figure 6).

**Figure 6.**
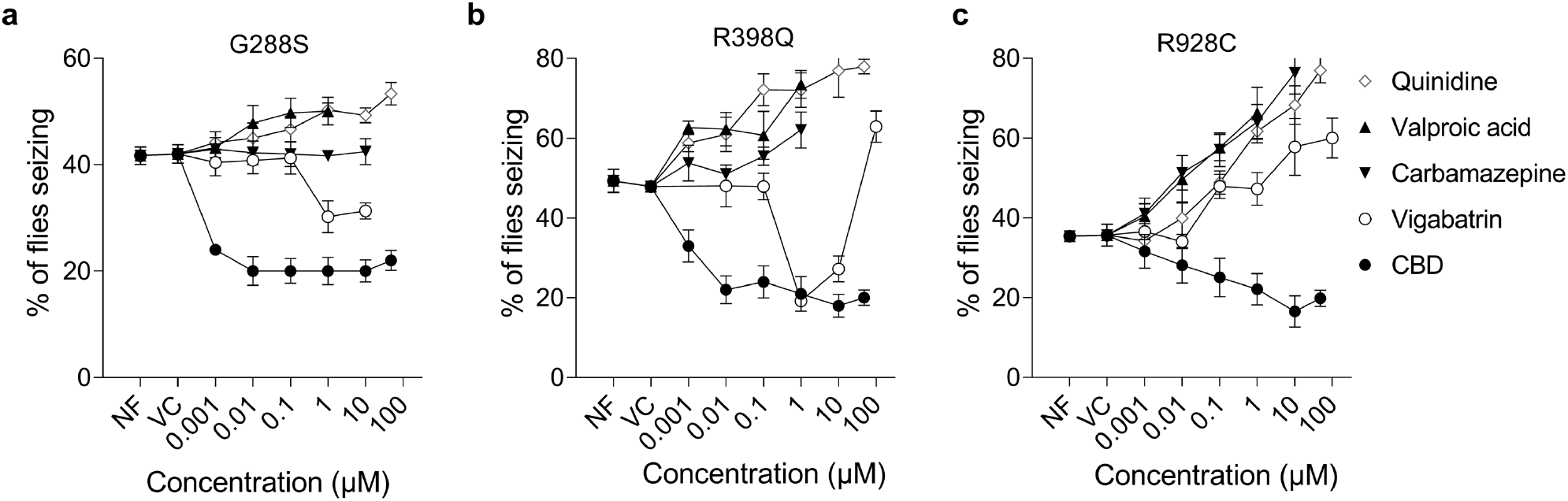
*In vivo* analysis of drug effects on seizure phenotype in *Drosophila* models. *Drosophila* expressing either WT or G288S, R398Q or R928C mutant human KCNT1 in GABAergic neurons were rasied from embryos on normal food (NF) or on food containing a range of concentrations of (**a**) CBD, (**b**) vigabatrin, (**c**) valproic acid, (**d**) carbamazepine or (**e**) quinidine and then analysed in the bang sensitive behavoural seizure assay. The percentage of *Drosophila* showing a seizure phenotype are shown for each dose of drug. All data points were compared to the vehicle control (VC) using a One-way ANAOVA with Dunnett’s multiple comparisons test. Compared to vehicle control, CBD significantly reduced seizures in G288S and R398Q mutants at all concentrations; and R928C at 10 and 50 μM. Vigabatrin significantly decreased seizures in G288S and R398Q mutants at 1 and 10 μM, but increased seizures at 100 μM in R398Q mutant and R928C mutant at 10 and 100 μM. Valproic acid had no effect on G288S mutant, but increased seizures in R398Q at 1 μM and R928C at 1 and 10 μM. Carbamazepine had no effect on G288S and R398Q mutants, but significantly increased seizures in R928C at 0.1, 1 and 10 μM. Quinidine exacerbated seizures in G288S mutant at 50 μM, and R398Q and R928C mutants at 0.1, 1, 10 and 50 μM. The exact P values for all data are presented in the Supplemental Table.

## Discussion

In this study we found that expression of human KCNT1 channels containing three patient-specific mutations gives a seizure phenotype in *Drosophila*. The phenotype was observed when the mutant channels were expressed in inhibitory GABAergic neurons. These results indicate that overactivity of the KCNT1 channel and increased K^+^ currents in inhibitory neurons is sufficient to induce a seizure phenotype in *Drosophila*. This finding is consistent with previous studies, including our recent analysis on the electrophsiological properties of a large series of patient *KCNT1* mutations (Rychkov et al., 2022). KCNT1 channels are integral membrane proteins thought to be important in depolarising the cell membrane, acting to inhibit excitability by reducing the likelihood of repetitive firing of action potentials. Greater silencing of inhibitory neurons by increased KCNT1 activity may lead to neuronal hyperexcitability, the mechanism that underlies seizures. Studies in mice also suggest that decreased excitability of inhibitory neurons contribute to seizures (Gertler et al., 2022; Shore et al., 2020). Expression of the three *KCNT1* mutants in all neurons, excitatory neurons or in glia in *Drosophila* were damaging, resulting in lethality. The reasons for this are not yet understood. The seizure phenotype of KCNT1 R398C was significantly stronger than that seen with R928C. This may be consistent with R928C being only identified in patients with the “milder” phenotype of sleep-related hypermotor epilepsy (SHE), while R398C (and G288S) are also found in patients with more severe phenotypes including epilepsy of infancy with migrating focal seizures (EIMFS)(Bonardi et al., 2021).

To ascertain whether our *KCNT1 Drosophila* models may be useful as tools for assessing drug treatmemts for *KCNT1*-epilepsy, we looked at the effects on the seizure phenotype for some of the drugs currently used to treat patients. We also looked at the *in vitro* effects of the drugs in cell models expressing KCNT1 channels to see if findings from the two systems are comparable. Feeding vigabatrin, valproic acid and carbamazeapine to *Drosophila* expressing wild type KCNT1 channels showed no evidence of toxicity or alteration of normal behaviour. In the *KCNT1* mutant lines, valproic acid and carbamazepine, both are known to inhibit voltage and use-dependent sodium channels (Ali et al., 2023), were seen to exacerbate the seizure phenotype in a dose-dependent manner. Vigabatrin, which increases GABA level by inhibiting GABA aminotransferase (Kobayashi, Endoh, Ohmori, & Akiyama, 2020), was seen to reduce the seizure phenotype at some doses in the animals expressing *KCNT1* G288S, and R398Q, and exacerbated the seizure phenotype in those expressing R928C. The seizure phenotype in the *Drosophila* models showed responses to the anti-epileptic drugs, with varied effects on the different mutant channels. Variable responses to the drugs are also seen in patients, with none being highly effective in inhibiting seizures. Further studies are needed to investigate if the *Drosophila* models will be useful in preclinical pharmacogenetics for predicting the response of patients with particular *KCNT1* mutations to different drugs.

Quinidine is a long known blocker of ion channels and has previously been used as an anti-arrhythmic drug (Paul, Witchel, & Hancox, 2002). It has previously been shown *in vitro* to significantly reduce the K^+^ currents of wild type (Yang et al., 2006) and epilepsy-associated mutant KCNT1 channels (Mikati et al., 2015; Milligan et al., 2014; Rizzo et al., 2016). However, it has shown variable results and serious side effects in some *KCNT1* epilepsy patients (Bearden et al., 2014; Bonardi et al., 2021; Chong, Nakamura, Saitsu, Matsumoto, & Kira, 2016; Fitzgerald et al., 2019; Gribkoff & Winquist, 2023; Mikati et al., 2015; Numis et al., 2018). Given the conflicting *in vitro* and *in vivo* (patients) results, we analysed the effects of quinidine on *Drosophila* expressing the G288S, R398Q or R928C *KCNT1* mutants. The R928C and R398Q mutant models each showed a strong dose-dependent increase in the seizure phenotype, which was less pronounced in the G288S mutant, and no effect was seen in animals expressing wild type *KCNT1*. The *in vivo* effects of quinidine on KCNT1 channels in this study therefore again differ from those seen in previous *in vitro* studies which show blocking of the channel activity. This is likely due to the presence, *in vivo*, of multiple interacting neural newtworks in the context of a functioning nervous system in a whole animal, and/or due to the inhibition of ion channels other than KCNT1 controlling the activity of the excitatory neurons, due to the higher concentrations of quinidine required to see the blocking effects in the *in vitro* studies (Gribkoff & Winquist, 2023).

Ingestion of medicinal cannabis derivatives containing the compound cannabidiol (CBD) reduces the frequency of seizures in some *KCNT1*-epilepsy patients and has not been reported to exacerbate seizures (Bonardi et al., 2021; Poisson et al., 2020). Although the exact mechanism of action of CBD in epilepsy treatment is yet to be elucidated, it has been postulated that CBD modulates intracellular calcium and adenosine mediated signalling to inhibit neuronal activity (Gray & Whalley, 2020; Tambe, Mali, Amin, & Oliveira, 2023). CBD was seen to significantly reduce the K^+^ currents in HEK293T cells expressing wild type and each of the G288S, R398Q, R298C mutant KCNT1 channels. The effect was rapid and stronger in ‘inside-out’ patches suggesting CBD may be acting directly on the cytoplasmic domain of the channel rather than through cannabinoid receptor signalling. Significantly, CBD was approximtaely 10 fold more efficient at blocking K^+^ currents of mutant KCNT1 channels than WT KCNT1 channels. While this preferential action on mutant KCNT1 channels is a highly desirable property for a *KCNT1* therapeutic, it is unclear why this might be the case. It is possible that CBD binding to KCNT1 channel is state-dependent and the higher open probability of the mutant KCNT1 channels (Rychkov et al 2022) results in a higher efficacy of the drug. While this, is the first demonstration of direct inhibition of the KCNT1 channel’s K^+^ currents by CBD further investigation into the mechanism of action on KCNT1 as well as the efficacy and dosing regimens for treatment of patients with CBD may also be warranted.

In summary, our study shows that the expression of patient-specific *KCNT1* mutations in *Drosophila* give a seizure phenotype, modelling human *KCNT1*-epilepsy. The seizure phenotypes in the *Drosophila* models were affected by the addition of drugs currently used to treat people with *KCNT1* epilepsy, suggesting they may be useful as preclinical tools for screening new therapeutics for these epilepsies. Importantly, we were able to demonstrate differential effects, some positive and others negative, as well as differential efficacies, of currently used drugs Together the three *KCNT1* mutations investigated in this study account for approximately one fifth of patients identified to date, thus dosing information and any candidate drugs identified using the models will potentially benefit a significant proportion of people with *KCNT1* epilepsy.

## Materials and Methods

### Generation of constructs and germ line transformation

Transgenic *Drosophila* were generated using the attP2 locus and PhiC31 integration system (Bischof, Maeda, Hediger, Karch, & Basler, 2007). Full-length cDNA for Human *KCNT1* Transcript 1 NM_020822.1 (Origene Technologies Inc, Rockville, MD, USA) was mutagenised to induce point mutations, G288S, R398Q and R928C using QuikChange lightning site-directed mutagenesis kit (Agilent Technologies Inc. Santa Clara, CA, USA).

Primers used for mutagenesis are as follows: p.G288S (5’-ACGGGGACCTGCAGCATCCAGCACC-3’ and 5’-GGTGCTGGATGCTGCAGGTCCCCGT-3’), p.R398Q (5’-CGCCCACCCCCAGCTCCAGGACT-3’ and 5’-AGTCCTGGAGCTGGGGGTGGGCG-3’) and p.R928C (5’-CCACCCTTCCAACATGTGCTTCATGCAGTTCCG-3’ and 5’-CGGAACTGCATGAAGCACATGTTGGAAGGGTGG-3’). The resulting constructs were cloned into EcoRI/XhoI restriction sites of pUAST-attB. All constructs were verified by Sanger DNA sequencing prior to injection into embryos (BestGene Inc. Chino Hills, CA, USA).

### Drosophila stocks and culture

*Drosophila* were cultured in 12-h light/dark cycles at specified temperatures on fortified medium (1% agar, 1% glucose, 6% fresh yeast, 9.3% molasses, 8.4% coarse semolina, 0.9% acid mix and 1.7% tegosept). The following *Drosophila* lines were obtained from the Bloomington *Drosophila* Stock Center (Indiana, USA): W118 (BL3605), *elav*^*C155*^*-GAL4* (BL458), *GAD1-GAL4* (BL51630), *Chat-GAL4* (BL56500) and *Repo-GAL4* (BL7415). The vector control BL8622 (y[1] w[67c23]; P{y[+t7.7]=CaryP}attP2) was used to generate transgenic *Drosophila* with human *KCNT1*.Transgenic *Drosophila* harbouring human *KCNT1* generated in this work were: *UAS-KCNT1 WT, UAS-KCNT1 G288S, UAS-KCNT1 R398Q* and *UAS-KCNT1 R928C*.

### Bang-sensitive behavioural assays to investigate a seizure phenotype

The bang-sensitivity behavioural assay (banging assay) was used to measure the recovery of *Drosophila* from seizure-like activities induced by mechanical shock and was performed as described previously (Tao et al., 2011). Experiments were performed between 8am and 11am to minimise the potential effects of circadian oscillation on animal activity. Between, ten and twenty *Drosophila* (females and males) aged between 4 to 8 days after eclosion were collected under CO_2_. The *Drosophila* were transferred to an empty clear 50ml measuring cylinder and allowed to acclimatise for 3 minutes before the cylinder was tapped on the bench to give twenty strong mechanical shocks. A Dino-Lite digital microscope (Product# AD3713TB; AnMo Electronics Corporation, New Taipei City, Taiwan) was used to record and save the videos of all of the experiments. Videos were replayed to score the *Drosophila* behaviour. *Drosophila* showing seizure-like behaviour were counted for 30 seconds. A minimum of 50 *Drosophila* were tested for each genotype. A one-way anova with Dunnett’s multiple comparisons test was used to determine the statistical difference of recovery from mechanical stimulation between different groups.

### Electrophysiology and analysis of effects of drugs

Whole-cell and inside-out patch-clamp recordings of KCNT1 mediated currents were performed using transiently transfected HEK293T cells 20-36 hr post transfection. Whole-cell patch clamping was performed using a computer-based patch-clamp amplifier (EPC-9, HEKA Elektronik) and PULSE software (HEKA Elektronik) as previously described (Rychkov et al., 2022). The bath solution contained 140 mM NaCl, 4 mM KCl, 2 mM CaCl2, 2 mM MgCl2 and 10 mM HEPES adjusted to pH 7.4 with NaOH. The pipette solution contained 80 mM K gluconate, 50 mM KCl, 10 mM NaCl, 1mM MgATP, 10 mM EGTA and 10 mM HEPES adjusted to pH 7.3 with KOH. Patch pipettes were pulled from borosilicate glass and fire polished to give a pipette resistance between 1 and 2 MΩ. Series resistance ranged between 2.5 - 4 MΩ and was 80-90% compensated. Only cells producing KCNT1 currents amenable to voltage clamp with the voltage error less than 10% due to a residual series resistance were used for the analysis of the drugs’ effects. In the inside-out patch clamp experiments pipettes with a resistance between 2 and 4 MΩ were filled with a standard bath solution (see above) and the membrane patch was perfused from the intracelular side with a solution containing 120 mM KCl, 20 mM NaCl, 0.2 mM EGTA and 10 mM HEPES adjusted to pH 7.3 with KOH. All drugs were dissolved in DMSO, aliquoted, and stored at -20C. The maximum DMSO concentration in the bath solution did not exceed 0.05%. Drugs were applied using a gravity-fed perfusion system with the outlet positioned within 1-2 mm of the patched cell and the perfusion rate of 0.5 ml/min.

## Supporting information

Supplementary Table of P vales for Figure 6

## Acknowledgments

L.M.D. was funded by an NHMRC Senior Research Fellowship 1104718, NHMRC Project Grant 1125523 and we thank the University of South Australia for funding assistance. C.X.L. was supported by a President’s Postgraduate Scholarship from the University of South Australia.

## Author Contributions

M.G.R. and L.M.D. designed and oversaw the study., R.H., C.X.L., A.I. and Z.S. generated constructs and performed the *Drosophila* experiments. E.A.C. and J.D.L. analysed and interpreted the electrophysiological experiments. G.Y.R. performed and interpreted the electrophysiological experiments. All authors wrote and edited the manuscript.

## Data availability statement

All data generated or analyzed during this study are included in this published article.

