## Supplementary Table of P vales for Figure 6 for "Modelling human KCNT1-epilepsy in *Drosophila*: a seizure phenotype and drug responses"

**Supplementary Table.** P values obtained using One way ANOVA with Dunnett’s multiple comparisons test of the data presented in Figure 6. Blue asterisks denote the statistically significant decrease in seizures compared to vehicle control, whereas red asterisks denote significant increase; ns – not significant. . N is the number of independent experiments; number in brackets is the total number of flies analysed in each condition. The number of the controls groups and the total number of flies for G288S, R398Q and R928C are 4 (54), 8 (314) and 4 (103) correspondingly.

|  | **G288S** | | | | | **R398Q** | | | | | **R928C** | | | | |
| --- | --- | --- | --- | --- | --- | --- | --- | --- | --- | --- | --- | --- | --- | --- | --- |
| **[µM]** | CBD | Vigabtr | Valpr | Carbam | Quin | CBD | Vigabtr | Valpr | Carbam | Quin | CBD | Vigabtr | Valpr | Carbam | Quin |
| **0.001** | <0.0001  ****  N=4(60) | 0.9604  ns  N=4(60) | 0.9999  ns  N=4(60) | 0.9996  ns  N=3(62) | 0.9979  ns  N=5(54) | 0.0136  *  N=4(73) | 0.1252  ns  N=5(101) | 0.0526  ns  N=5(54) | 0.6311  ns  N=5(77) | 0.3140  ns  N=4(70) | 0.9608  ns  N=4(73) | 0.9998  ns  N=5(70) | 0.8910  ns  N=5(54) | 0.8675  ns  N=3(50) | 0.9996  ns  N=4(67) |
| **0.01** | <0.0001  ****  N=4(60) | 0.9819  ns  N=4(60) | 0.4985  ns  N=4(60) | 0.9977  ns  N=3(60) | 0.9758  ns  N=5(50) | <0.0001  ****  N=4(50) | >0.9999  ns  N=4(50) | 0.0613  ns  N=5(50) | 0.9479  ns  N=5(74) | 0.1634  ns  N=4(66) | 0.5964  ns  N=4(52) | 0.9996  ns  N=5(79) | 0.0904  ns  N=5(50) | 0.0550  ns  N=4(54) | 0.8641  ns  N=4(51) |
| **0.1** | <0.0001  ****  N=4(60) | 0.9944  ns  N=4(60) | 0.2154  ns  N=4(60) | 0.9924  ns  N=4(55) | 0.7269  ns  N=5(50) | <0.0001  ****  N=6(158) | >0.9999  ns  N=4(76) | 0.1073  ns  N=5(50) | 0.4074  ns  N=5(51) | 0.0012  **  N=5(54) | 0.1644  ns  N=6(94) | 0.2033  ns  N=4(74) | 0.0037  **  N=5(50) | 0.0023  **  N=5(55) | 0.0125  *  N=6(58) |
| **1** | <0.0001  ****  N=4(60) | 0.0124  *  N=4(60) | 0.1815  ns  N=4(60) | 0.9997  ns  N=3(52) | 0.1240  ns  N=4(55) | <0.0001  ****  N=5(126) | <0.0001  ****  N=5(132) | 0.0006  ***  N=5(68) | 0.0569  ns  N=6(58) | 0.0022  **  N=4(55) | 0.0583  ns  N=5(91) | 0.1963  **  N=5(99) | <0.0001  ****  N=5(50) | 0.0003  ***  N=3(53) | <0.0001  ****  N=5(51) |
| **10** | <0.0001  ****  N=4(60) | 0.0240  *  N=4(60) | 0.9999  ns  N=4(60) | 0.9996  ns  N=3(65) | 0.2295  ns  N=5(50) | <0.0001  ****  N=6(117) | 0.0010  **  N=4(106) |  |  | 0.0001  ***  N=5(66) | 0.0021  **  N=6(94) | 0.0014  *** N=5(80) |  | <0.0001  ****  N=5(67) | <0.0001  ****  N=5(51) |
| **50** | <0.0001  ****  N=4(60) |  |  |  | 0.0153  *  N=5(50) | <0.0001  ****  N=7(96) |  |  |  | <0.0001  ****  N=5(59) | 0.0096  ** N=7(86) |  |  |  | <0.0001  ****  N=5(56) |
| **100** |  |  |  |  |  |  | 0.0153  *  N=5(62) |  |  |  |  | 0.0005  ***  N=5(75) |  |  |  |
